# Enhanced synaptic properties of the prefrontal cortex and hippocampus after learning a spatial working memory task in adult male mice

**DOI:** 10.1101/339432

**Authors:** Vasiliki Stavroulaki, Vasileios Ioakeimidis, Xanthippi Konstantoudaki, Kyriaki Sidiropoulou

## Abstract

Working memory (WM) is the ability to hold on-line and manipulate information. The prefrontal cortex (PFC) is a key brain region involved in WM, while the hippocampus is also involved, particularly, in spatial WM. Although several studies have investigated the neuronal substrates of WM in trained animals, the effects and the mechanisms underlying learning WM tasks have not been explored. In our study, we investigated the effects of learning WM tasks in mice on the function of PFC and hippocampus, by training mice in the delayed alternation task for 9 days (adaptive group). This group was compared to naïve mice that stayed in their homecage (naïve) and mice trained in the alternation procedure only (non-adaptive). Following training, a cohort of mice (Experiment A) was tested in the left-right discrimination task and the reversal learning task, while another cohort (Experiment B) was tested in the attention set-shifting task (AST). The adaptive group performed significantly better in the reversal learning task (Experiment A) and AST (Experiment B), compared to non-adaptive and naïve groups. At the end of the behavioral experiments in Experiment A, field excitatory post-synaptic potential (fEPSP) recordings were performed in PFC and hippocampal brain slices. The adaptive group had enhanced the long-term potentiation (LTP) in the PFC, compared to the other groups. In the hippocampus, both the adaptive and the non-adaptive groups exhibited increased fEPSP compared to the naive group, but no differences in LTP. In Experiment B, the dendritic spine density was measured, which, in the PFC, was found increased in the adaptive group, compared to the non-adaptive and naive groups. In the hippocampus, there was an increase in mature dendritic spine density in the adaptive group, compared to the other two groups. Our results indicate a role for long-term potentiation and dendritic spine density in learning WM tasks.

**Significance statement:** Working memory (WM) allows for transient storage and manipulation of information and has a central role in cognition. While a great number of research studies have investigated the mechanisms underlying the ‘memory’ part of WM in well-trained animals, the mechanisms that underlie learning WM tasks are not known. Studies have indicated that learning a WM tasks alters and enhances neuronal firing during the delay period, suggesting that long-term plasticity mechanisms could be involved. Our results in this study suggest that learning a working memory task primarily increases long-term potentiation and dendritic spine density in the prefrontal cortex, providing evidence for a role of long-term plasticity processes in learning working memory tasks. Furthermore, learning working memory tasks enhances cognitive flexibility.

## 1. Introduction

Working memory (WM) refers to the ability to maintain and manipulate information on- line (Goldman-Rakic, 1995). WM is supported by neuronal networks across various brain regions, including the prefrontal cortex (PFC), the parietal and temporal cortical regions as well as the hippocampus (Constantinidis and Procyk, 2004; Rissman et al., 2007; Spellman et al., 2015; Boku et al., 2017; Tamura et al., 2017). The main cellular mechanisms proposed to underlie WM function is persistent activity during the delay period of WM tasks in primates (Riley and Constantinidis, 2015), which has also been observed in rodent studies (Baeg et al., 2003; Liu et al., 2014; Kamigaki and Dan, 2017). However, different patterns of neuronal activity have been recorded during the delay period (Narayanan and Laubach, 2009), indicating that neuronal activity does extend throughout the delay period. Thus, other mechanisms have been suggested to underlie WM function, including short-term plasticity (Mongillo et al., 2008) and neuronal synchronization (Lundqvist et al., 2016, 2018).

When PFC neurons were recorded throughout the learning period of WM tasks, two main changes have been identified in both primates and rodents: a) more neurons exhibit persistent neuronal activity during the delay period and b) the firing frequency of persistent neuronal activity during the delay period is increased (Baeg et al., 2003; Meyer et al., 2007; Liu et al., 2014), indicating a role of long-term plasticity process in learning WM tasks. However, there are few studies implicating the role of long-term plasticity processes in WM, which have shown a correlation between long-term potentiation in the PFC and learning rate of WM tasks (Konstantoudaki et al., 2017; Chalkiadaki et al., 2019). Dendritic spines are protrusions located on dendritic branches of neurons and support all the different plasticity process of excitatory synapses (Sala and Segal, 2014; Lisman, 2017) and also support WM and spatial tasks (Brennaman et al., 2011; Velázquez-Zamora et al., 2011, 2012; Mahmmoud et al., 2015).

In our study, we trained mice in the delayed alternation task and then investigated long-term potentiation in PFC and hippocampal brain slices, as well as dendritic spine density. This methodology was chosen because training in WM tasks has been suggested to improve other cognitive functions, in particular cognitive flexibility (Light et al., 2010). Therefore, we examined cognitive flexibility using the reversal learning and the attention set-shifting tasks in the mice that were trained in WM tasks. We included two different types of controls: a) a passive control group, which stayed in the homecage and b) an active control group which performed an alternation task without any delays.

## 2. Materials and Methods

### 2.1 Animals

All mice were bred and housed in the IMBB-FORTH facility. Mice were housed in groups (3–4 per cage) and provided with standard mouse chow and water ad libitum, under a 12 h light/dark cycle (light on at 8:00 a.m.) with controlled temperature (24 +/− 1°C). All procedures were performed according to protocols approved by the Research Ethics Committee of the University of Crete and obey the European Union ethical standards outlined in the Council Directive 2010/63 EU of the European Parliament on the protection of animals used for scientific purposes.

Two different groups of male mice were used for the experiments: a) F1 generation of 129/SVX C57/B6 mice, aged 7-8 months old and b) C57/B6 mice aged 7-8 months old. We chose this age because WM is one of the first to decline during the normal aging process (Bucur and Madden, 2010; Wang et al., 2011) along with alternations in delay activity (Wang et al., 2011), in neuronal morphology and in excitability (Samson and Barnes, 2013). Young adult mice (3-4 months) from both strains were also used for some experiments.

### 2.2 Behavioral tasks

The T-maze apparatus used includes a start arm and 11 two goal arms (45×5cm each). The left-right discrimination task used examines reference memory in mice (Shoji et al., 2012), while reversal learning is a form of cognitive flexibility (Bissonette and Powell, 2012). The delayed alternation task is a classic task used for the study of working memory and was performed as described before (Konstantoudaki et al., 2017). In both experiments, mice were trained in the delayed alternation task. Mice were initially handled by the experimenter for 10 days, and then habituated in the T-maze apparatus, for 2 days. Mice were food-restricted so that the animals maintained 85-90% of their initial weight. During the second habituation day, the time that each mouse spent in each arm was calculated in order to establish the arm preference for each mouse.

In experiment A (Figure 1A), mice were initially trained in the left-right discrimination (LRD) task for 2 days. For each mouse, the arm that was baited was the non-preferred arm for that mouse, based on the measurements during the second habituation day. Each mouse, individually, was subjected to a single 20-trial session each day. After each trial, the mouse was placed in a cage for 1 min. On the second day, following the 20-trial session, reversal learning was examined by placing the reward in the opposite arm of the one that was during the LRD task for 10 trials. Mice were then split into three groups. Mice in the naive group sat in their homecage, while mice in non-adaptive and adaptive groups continued training in the T-maze, this time in the alternation task. All the mice were subject to 10-trial sessions, 3 sessions/day. At the first trial of each session, mice were allowed to freely choose between the right or left goal arms. In the following trials, mice had to alternate the goal arms in order to receive reward, initially with no temporal delay between the trials. Once they reached a predefined criterion for the alternation procedure [i.e., 2 consecutive sessions of ≥70% correct choices (performance)], mice were split into the non-adaptive and adaptive groups. Mice in non-adaptive group continued to perform the same alternation task for 2 sessions per day. Instead, mice in the adaptive group started the delayed alternation procedure, for which delays were introduced, initially at 10 seconds and increasing by 10 seconds when the criterion for each delay was achieved, for 9 days. After completion of the delayed alternation procedure, mice were asked to ‘remember’ the left-right discrimination task, for 20 sessions, and to adjust to reversal of the reward, for another 10 sessions.

**Figure 1.**
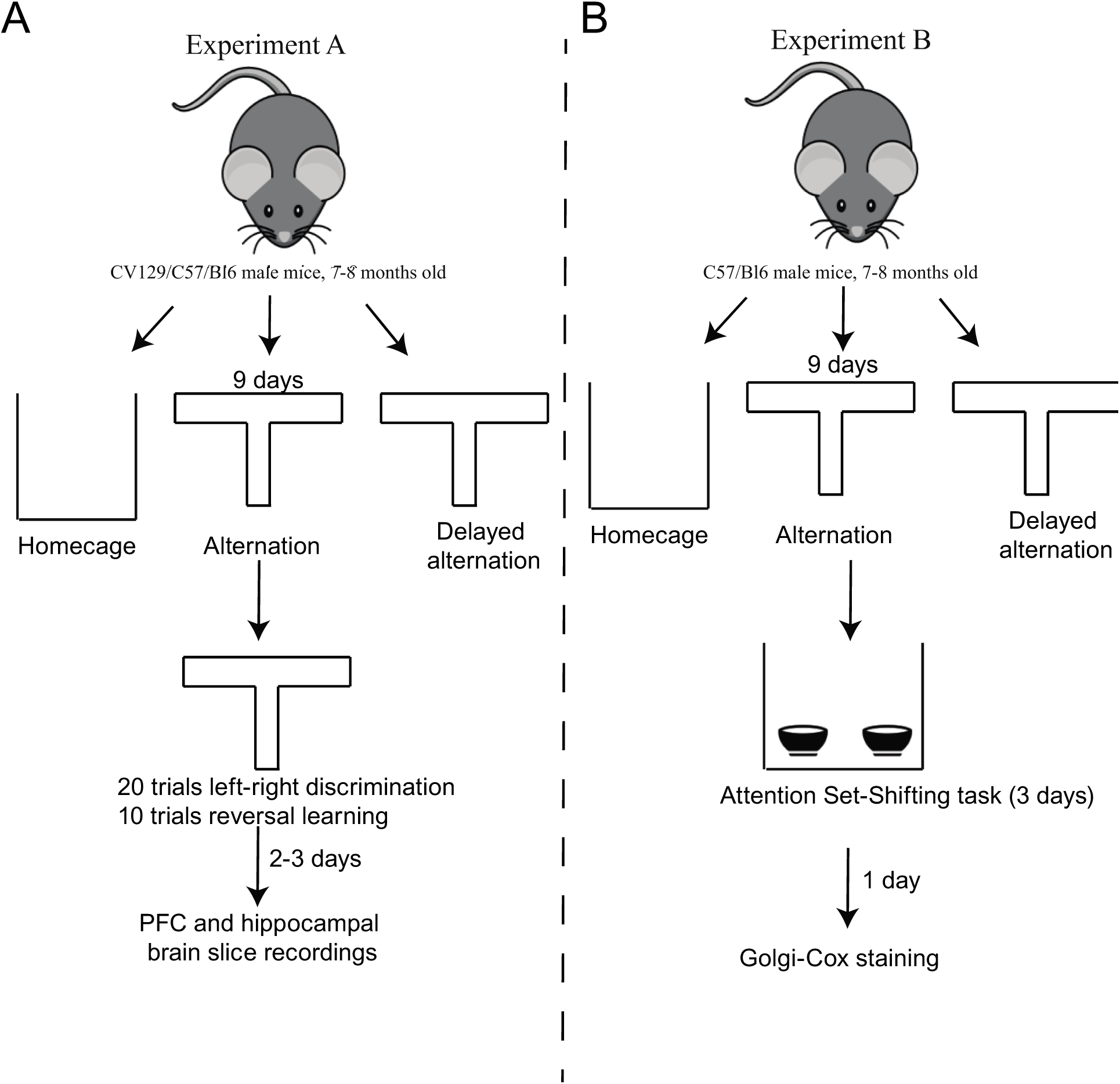
Experimental design of the study, outlining the two different experiments (A and B), the three different groups used, namely, the naïve (homecage), the non-adaptive (alternation procedure) and the adaptive (delayed alternation procedure) groups and the different tasks used in each experiment. A. In experiment A, mice were trained for 9 days and, then, tested in the left-right discrimination task and the reversal task, followed by brain slice electrophysiological experiments to study the evoked fEPSP and LTP in the PFC and the hippocampus. B. In experiment B, mice were trained for 9 days and, then, tested in the AST task, followed by Golgi-Cox staining and measurements of dendritic spine density in the PFC and the hippocampus.

In Experiment B, (C57/Bl6 mice) were split into three groups and trained accordingly, as described above (Figure 1B). Following training, all groups of mice were subjected to the attentional set shifting task (AST), a behavioral task that examines cognitive flexibility (Brown and Tait, 2016). For this task, an open field device is used along with two small identical cups, which contain various substrates, the olfactory cues and food reward. For 2 days, mice habituate to the open-field and they learn to dig for food in the bowl filled with sawdust. During the different experimental stages, mice need to pay attention and respond to the relevant cue (digging medium) and ignore an irrelevant cue (odor), by pairing a food reward with the medium. This association is then reinforced in subsequent stages where the type of digging medium and odor changes, before changed entirely in the final stage. This task contains 7 stages, namely: a) simple discrimination (SD) in which mice differentiate between two different beddings, b) compound discrimination (CD), in which mice still differentiate between two different beddings but two different olfactory cues are introduced that are not relevant, c) compound discrimination reversal (CDR) in which the bedding and olfactory cues are the same, but the rewarded bedding is switched, d) intradimentional shift 1 (ID-1), in which two different bedding media and olfactory cues are introduced, but one of the beddings is rewarded, e) intradimentional shift 2 (ID-2), in which two different bedding media and olfactory cues are introduced, but one of the beddings is rewarded, f) intradimentional shift 2 reversal (ID-2R), in which the bedding and olfactory cues are the same as in ID-1 but the rewarded bedding is switched, and g) extradimentional shift (ED), in which one of the olfactory cues is rewarded. For mice to successfully completie each stage, they need to complete six consecutive correct trials. The total # of trials required to reach criterion were recorded for each stage.

### 2.3 Electrophysiological recordings

In Experiment A, 1-3 days following the end of the last behavioral session, mice were prepared for electrophysiological experiment, using in vitro slice preparation. This range was necessary to be able to test all mice in the behavioral and the electrophysiological experiments. We did not observe any difference that was dependent on the day of the recording. The person performing the electrophysiological experiments was blind to the type of behavioral training on the animals. Mice were decapitated under halothane anesthesia. The brain was removed immediately and placed in ice cold, oxygenated (95% O_2_/5% CO_2_) artificial cerebrospinal fluid (aCSF) containing (in mM): 125 NaCl, 3.5 KCl, 26 NaHCO_3_, 1 MgCl_2_ and 10 glucose (pH=7.4, 315 mOsm/l). The brain was blocked and glued onto the stage of a vibratome (Leica, VT1000S, Leica Biosystems GmbH, Wetzlar, Germany). Brain slices (400μm thick) containing either the PFC or the hippocampus were taken and were transferred to a submerged chamber, which was continuously superfused with oxygenated (95% O2/5% CO2) aCSF containing (mM): 125 NaCl, 3.5 KCl, 26 NaHCO_3_, 2 CaCl_2_, 1 MgCl_2_ and 10 glucose (pH=7.4, 315mOsm/l) in room temperature. The slices were allowed to equilibrate for at least an hour in this chamber before experiments began. Slices were then transferred to a submerged recording chamber, which continuously superfused oxygenated (95% O2/5% CO2) aCSF containing (in mM): 125 NaCl, 3.5 KCl, 26 NaHCO_3_, 2 CaCl_2_, 1 MgCl_2_ and 10 glucose (pH=7.4, 315mOsm/l) in room temperature.

The extracellular recording electrode, filled with NaCl (2M), was placed within the upper layers of the PFC or BC. The platinum/iridium metal microelectrode (Harvard apparatus UK, Cambridge, UK) was also placed within the upper layers of the PFC or the stratum radiatum layer of the CA1 region of the hippocampus, about 300μm away from the recording electrode, and was used in order to evoke fEPSPs. Responses were amplified using a Dagan BVC-700A amplifier (Dagan Corporation, Minneapolis, MN, USA), digitized using the ITC- 18 board (Instrutech, Inc) on a PC using custom-made procedures in IgorPro (Wavemetrics, Inc, Lake Oswego, OR, USA). Data were acquired and analyzed using custom-written procedures in IgorPro software (Wavemetrics, Inc, Lake Oswego, OR, USA).

The electrical stimulus consisted of a single square waveform of 100 μsec duration given at intensities of 0.05-0.3 mA generated by a stimulator equipped with a stimulus isolation unit (World Precision Instruments, Inc). The fEPSP amplitude was measured from the minimum value of the synaptic response (4-5 ms following stimulation) compared to the baseline value prior to stimulation. Both parameters were monitored in real-time in every experiment. A stimulus-response curve was then determined using stimulation intensities between 0.05-0.3 mA. For each different intensity level, two traces were acquired and averaged. Baseline stimulation parameters were selected to evoke a response of 1mV. The paired-pulse protocol consisted of two pulses at baseline intensity separated by 100, 50 and/or 20 msec. For the LTP experiments in the PFC, baseline responses were acquired for at least 20 minutes, then three 1second tetanic stimuli (100Hz) with an inter-stimulus interval of 20 seconds were applied and finally responses were acquired for at least 50 minutes post-tetanus. For the experiments in the hippocampus, the recording and stimulating electrodes were placed on the stratum radiatum of the CA1 region, about 300μm apart. LTP was induced using theta-burst stimulation, which consisted of 5 pulse at 100Hz, repeated four times at theta-rhythm (every 200ms). This stimulation was repeated twice with an inter-stimulus interval of 20 seconds. Synaptic responses were normalized to the average 10 minutes pre-stimulus (tetanus or theta-burst).

### 2.4 Golgi Cox staining

After the AST task the mice in the second cohort were euthanised and their brains were removed and subjected to Golgi Cox staining. This staining remains a key method to study neuronal morphology (Zaqout and Kaindl, 2016). Specifically, mice brains were removed, isolated and placed in a glass vial in a Golgi Cox solution (5% Potassium Dichromate in dH2O, 5% Mercuric Chromate in dH2O, 5% Potassium Chromate in dH2O), that it made and stored 5 days in the dark. The solution is renewed day by day for 10 days. The eleventh day, brains transfer to a 30% sucrose solution in dH2O at 4° C, until the day they were slicing with a vibratome. 150-μm coronal slides of the prefrontal cortex and hippocampus were obtained with a vibratome (VT1000S, Leica Microsystems, Wetzlar, Germany) and placed on gelatin-coated slides. After finishing the sectioning of all samples, the slices kept at 4°C in dark for one day.

For staining processing, the tissue sections were treated with ammonium hydroxide and fixed with Kodak rapid fixer (30 min per solution) following the procedure described by Gibb and Kolb (1998). Finally, the tissue was rinsed and dehydrated in graded concentrations of alcohol and xylene. At the end, the slides mounting and covered with cover glass.

One month later, the slices were observed and photographed under a light microscope (Nikon Plan Apo 60x / 140oil WD 0.21 lenses with a Nikon FDX-35 camera). Stacks were created from images of each cell of pyramidal secondary dendrites of PFC and HPC in order to measure the number, length and density of spines. All images are processed using Adobe Photoshop 7.0 and ImageJ Software and Cell counter. The spines were subdivided into mushroom / thin (referred as mature) and stubby groups.

### 2.4 Statistical Analysis

Data analysis was performed with Microsoft Excel. The data were first tested for normality. One-way, repeated measures ANOVA or t-tests were performed depending on the experiment. Statistical analysis was performed in IBM SPSS Statistics v.22. Data are presented as mean ± standard error of mean (SEM).

## 3. Results

### 3.1 Experiment A

In the first experiment, thirty-one adult male mice, 129/SV-C57/B6 were used. Mice were first trained to acquire the LRD task in the T-maze and were exposed to 10 trials of reversal learning. They subsequently split into three groups (Figure 1): a) the naïve group, in which the mice stayed in their home-cage (passive control), n=9, b) the non-adaptive group in which mice underwent training the alternation task in the T-maze, without the introduction of delays, a group that served as the active control group, n=11, and c) the adaptive group, in which mice underwent delayed alternation task were trained in a spatial working memory task, n=11 namely the delayed alternation in the T-maze.

Mice of all three groups learned the LRD task equally well. There were no differences between the three groups in both days of training required to reach the criterion of learning the task (one-way ANOVA, F_(2,29)_=0.25, p=0.5) (Figure 2A). Similarly, there was no difference in the performance of the reversal learning task in all three groups of mice, prior to working memory training (one-way ANOVA, F=0.12, p=0.6) (Figure 2D).

**Figure 2.**
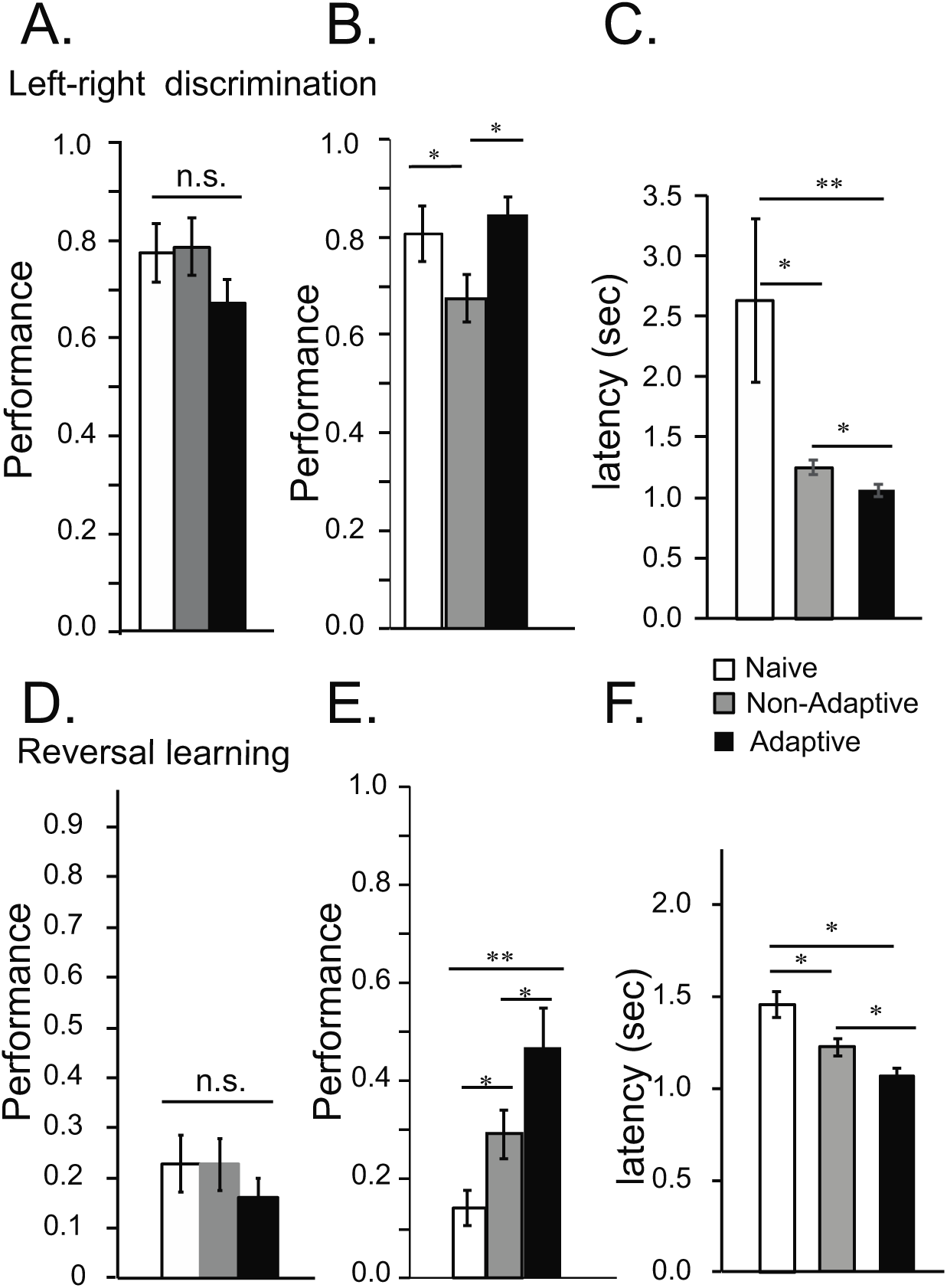
Performance of mice in the left-right discrimination and reversal tasks before and after the delayed alternation task. A. Bar graphs showing percent correct in the first and second days of reference memory acquisition in the left-right discrimination task, before WM training. No significant difference among the three groups was identified (one-way ANOVA, F_(2,29)_=0.25, p=0.5). B. Bar graphs showing percent correct in the left-right discrimination task after WM training. Significant difference was identified among the three groups (one-way ANOVA, F_(2,29)_=2.4, p=0.04). Post-hoc Tukey comparisons identified significant difference between the non- adaptive and the adaptive groups (Tukey test, p=0.03) and not between the non-adaptive and the naïve groups (Tukey test, p=0.3) or between the naïve and the non-adaptive groups (Tukey test, p=0.15). C. Bar graphs showing the latency (sec) in the left-right discrimination task after WM training. Significant difference was identified among the three groups (one-way ANOVA, F_(2,29)_=7.3, p=0.003). Post-hoc comparisons identified significant differences between the non-adaptive and the naive groups (Tukey test, p=0.001) as well as between the adaptive and naïve groups (Tukey test, p=0.01), and between the non-adaptive groups and adaptive groups (Tukey test, p=0.03). D. Bar graphs showing percent correct in the reversal learning task, before WM training. No significant difference among the three groups was identified (one-way ANOVA, F_(2,29)_=0.12, p=0.5) E. Bar graphs showing percent correct in the reversal learning task after WM training. Significant difference was identified among the three groups (one-way ANOVA, F_(2,29)_=5.5, p=0.001). Post-hoc comparisons identified significant differences between the non-adaptive and the adaptive groups (Tukey test, p=0.02) as well as between the adaptive and naïve groups (Tukey test, p=0.007), but not between the naïve and the non-adaptive groups (Tukey test, p=0.8). F. Bar graphs showing the latency (sec) in the reversal learning task after WMT. Significant difference was identified among the three groups (one-way ANOVA, F_(2,29)_=10.3, p=0.001). Post-hoc comparisons identified significant differences between the non-adaptive and the naive groups (Tukey test, p=0.03) as well as between the adaptive and naïve groups (Tukey test, p=0.001), and between the non-adaptive groups and adaptive groups (Tukey test, p=0.04).

Following training in the delayed alternation task, mice were tested again in the LRD and the reversal learning tasks in the T-maze. There was a significant difference among the three groups in the performance of the LRD task (one-way ANOVA, F_(2,29)_=2.4, p=0.04) (Figure 2B). Post-hoc analyses revealed that the adaptive group performed significantly better compared to mice in the non-adaptive group, but equally well compared to naïve in the LRD task (p=0.3) (Figure 2C). A significant effect of training was also found in the reversal learning task (one-way ANOVA, F_(2,29)_=5.5, p=0.001). Specifically, mice in the adaptive group performed significantly better compared to mice in the non-adaptive and the naïve groups in the reversal learning task (Tukey test, p=0.02 and p=0.007, respectively, Figure 2E). There was no significant difference between the naive and the non-adaptive groups in the reversal learning task (p=0.4). In addition, the latency was significantly decreased in the both the adaptive and non-adaptive groups, compared to the naïve group, in the LRD task, while latency was reduced in the adaptive group, compared to both the non-adaptive and naïve groups, in the reversal learning task (Figure 2C, F).

Following the behavioral testing, synaptic transmission and synaptic plasticity in the PFC and the hippocampus were studied, using the brain slice preparation. Field EPSPs were recorded in PFC layer II while stimulating layer II. There was no significant difference in the fEPSP recorded in response to increasing stimulation in the PFC between the naive, the non-adaptive and the adaptive groups (repeated measures ANOVA, F_(2,19)_=0.7, p=0.5) (Figure 3A). On the other hand, there was a significant difference in the fEPSPs recorded in the CA1 region while stimulating the Schaffer collateral axons (repeated-measures ANOVA, F_(2,19)_=5.5, p=0.01). Specifically, the fEPSPs in both the non-adaptive and adaptive groups of mice were increased, compared to the naive group (Tukey test, p=0.01 for the adaptive and p=0.02 for the non-adaptive group) (Figure 3B). This suggests that training in a spatial WM task, namely the alternation task, with delays or not, increases the efficacy of synaptic transmission in the CA1 region of HPC.

**Figure 3.**
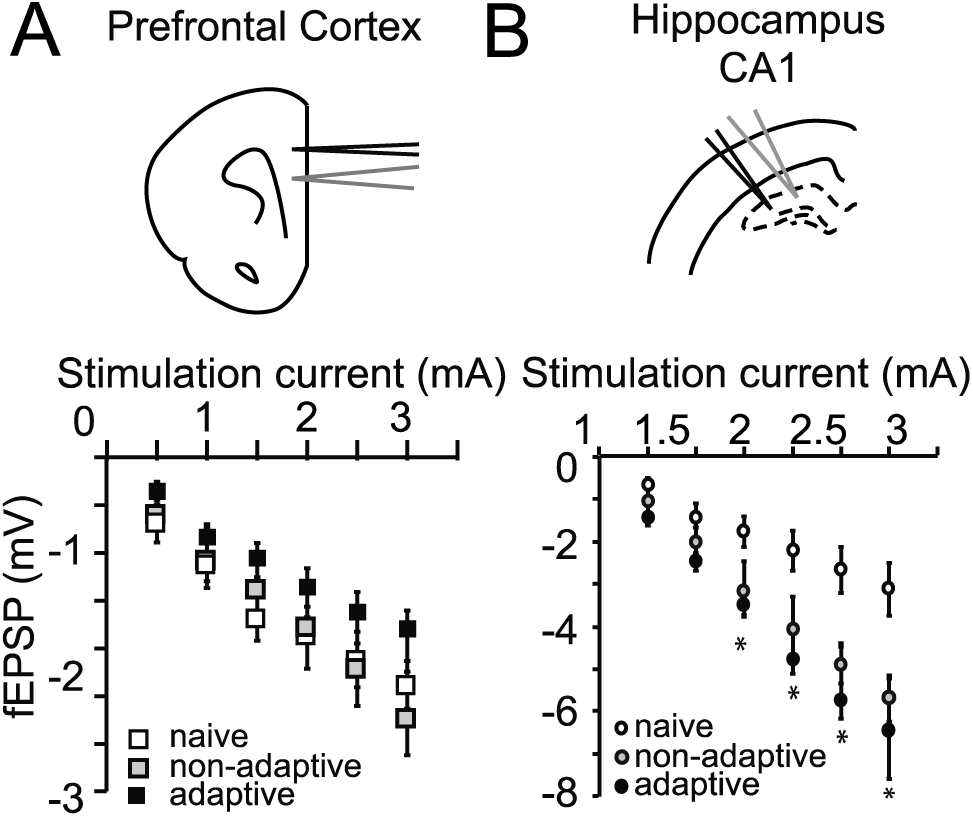
Synaptic transmission in the PFC and the hippocampus, in the naïve, non-adaptive and adaptive groups. A. Electrode layout in the PFC. B. Graph showing that there is no difference in the fEPSP amplitude in response to increasing current stimulation in the PFC (repeated measures ANOVA, F_(2,19)_=0.7, p=0.5). C. Electrode layout in the hippocampus. D. Graph showing that there is a significant difference in the fEPSP amplitude in the response to increasing current stimulation in the hippocampus, among the three different groups (repeated measures ANOVA, F_(2,19)_=5.5, p=0.01). Post-hoc comparisons show significant differences between the non-adaptive and naïve groups, as well as between the adaptive and naïve groups.

Next, we examined the induction and maintenance of LTP in both the PFC and hippocampus of mice from the three training groups. We find that tetanic stimulation resulted in a small, non-significant potentiation of the fEPSP in mice of the naïve and non-adaptive groups. This suggests that LTP in the middle-aged PFC has significantly decreased compared to the early-adulthood (3-5 months old) (repeated measures ANOVA, F_(1,15)_=8.2, p=0.01, Figure 4A), when the fEPSP is increased following tetanic stimulation as we have shown in our previous studies (Konstantoudaki et al., 2017; Chalkiadaki et al., 2019). Among the three different training groups, there was a significant difference in the fEPSP potentiation (repeated-measures ANOVA, F_(2,19)_=4.3, p=0.03, Figure 4B). Post-hoc comparisons revealed that the fEPSP potentiation was significantly greater in the adaptive group, compared to the non-adaptive and naïve groups, 30-50min after the tetanic stimulation (Tukey’s test, p=0.03). These results suggest that, while LTP has been reduced to non-significant levels at 7 months of age in mice, training in a WM task allows for re-emergence of LTP in PFC layer II synapses of middle-aged mice. In the hippocampus, theta-burst stimulation resulted in fEPSP potentiation in all groups of mice, naïve, non-adaptive and adaptive (Figure 4C). There was no significant difference in the LTP between the three groups (repeated measures ANOVA, F_(2,19)_=1.6, p=0.2).

**Figure 4.**
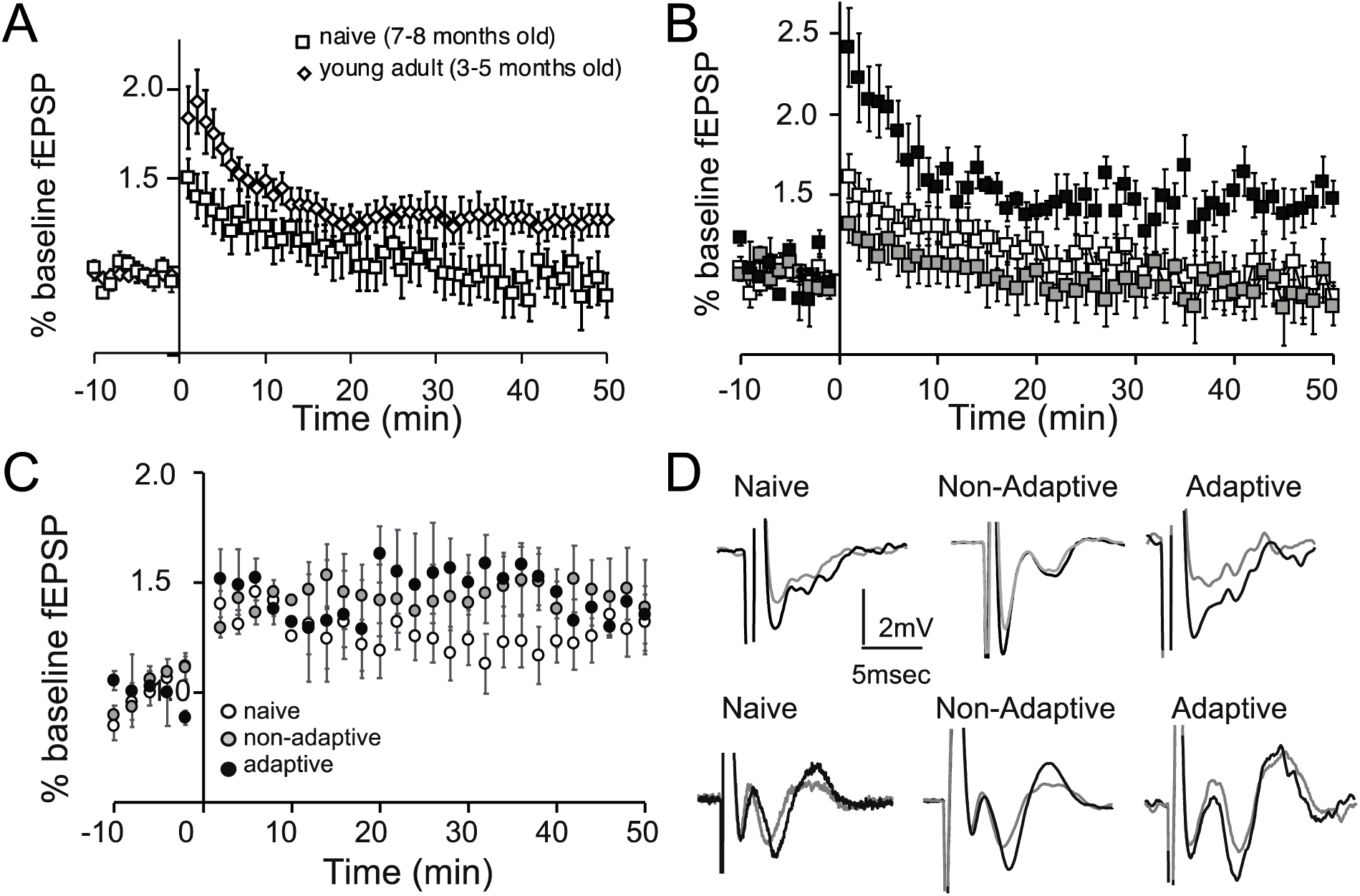
Long-term potentiation in the PFC and the hippocampus of naïve, non-adaptive and adaptive groups. A. Graph showing the potentiation of the fEPSP following tetanic stimulation in the PFC of naïve (7-8 month old) mice and naïve young adult mice (3-5 month old) (repeated measures ANOVA, F_(1,15)_=8.2, p=0.01) B. Graph showing the potentiation of the fEPSP following tetanic stimulation in the PFC of the three different training groups. There was a significant difference between the three groups (repeated measures ANOVA, F_(2,19)_=4.3 p=0.03). Post-hoc comparisons showed that there is a significant difference between the adaptive and naïve (Tukey test, p=0.02) and the adaptive and non-adaptive groups (Tukey test, p=0.03) for the time points 10min to 50min. C. Graph showing the potentiation of the fEPSP following theta-burst stimulation in the hippocampus. There was no significant difference between the three groups (repeated measures ANOVA, F_(2,19)_=1.6, p=0.2) D. Representative traces of the PFC (upper traces) and hippocampal (lower traces) LTP in the three different training groups.

### 3.2 Experiment B

In the second experiment, 25 C57/Bl6 male mice were used, which were also split into 3 groups: a) naïve (n=7), b) non-adaptive (n=9) and c) adaptive (n=9) groups. Mice in the naïve group stayed in their homecage for 9 days, mice in the non-adaptive group underwent the alternation task for 9 days, while mice in the adaptive group underwent the delayed alternation task for 9 days. Following training, mice of all three groups were subjected to the AST task (Figure 1B). One-way ANOVA analyses revealed non-significant effects of training on the SD (F_(2,22)_=0.18, p=0.19), CD (F_(2,22)_=0.03, p=0.97), ID-1 (F_(2,22)_=0.74, p=0.48), ID-2 (F_(2,22)_=2.6, p=0.1), ID2-R (F_(2,22)_=1.6, p=0.2) parts of the task. On the other hand, there was a significant effect of training in the CDR (one-way ANOVA, F_(2,22)_=5.5, p=0.01) and the ED (one-way ANOVA, F_(2,22)_=4.4, p=0.02) parts of the task (Figure 5). Post-hoc analyses showed that in the CDR part, adaptive mice required reduced number of trials to reach criterion compared to the non-adaptive mice (Tukey’s test, p=0.01). In the ED part of the task, post-hoc analyses showed that the adaptive group performed significantly better compared to the non-adaptive and naïve groups (Tukey’s test, p=0.04 and p=0.03, respectively). These results show that WM training improves performance in another test of cognitive flexibility, in addition to the reversal learning shown before.

**Figure 5.**
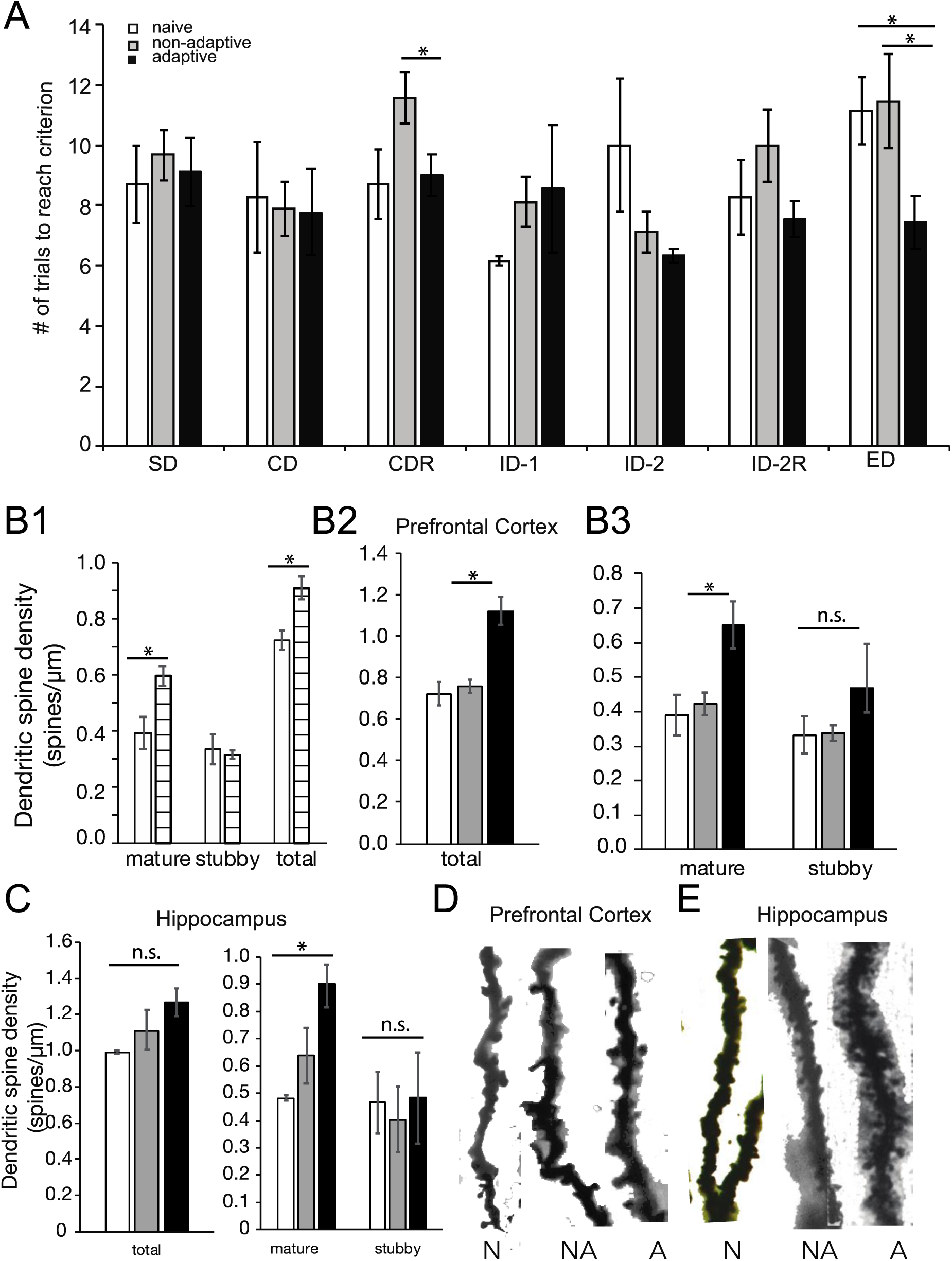
Performance in the AST task and dendritic spine density of naïve, non-adaptive and adaptive mice. A. Graph showing the average total number of trials that mice of the three different groups had to complete to reach criterion and successfully complete each part of the task. SD: simple discrimination, CD: compound discrimination, CDR: compound discrimination reversal, ID- 1: intradimentional shift-1, ID-2: intradimentional shift-2, ID-2R: intradimentional shift-2 reversal, ED: extradimentional shift. Significant differences among the three groups were identified in the CDR and the ED parts of the task (one-way ANOVA, F_(2,22)_=5.5, p=0.01 and F_(2,22)_=4.4, p=0.02, respectively). B1. Graph showing the mature, stubby and total dendritic spine density in the PFC of naïve (7- 8 month old) mice, compared to young adult mice. A significant decrease is found in the mature and total dendritic spine density (t-test, p=0.03). B2. Graph showing the total dendritic spine density in the PFC of naïve, non-adaptive and adaptive groups of mice. B3. Graph showing the mature and stubby dendritic spine density in the PFC of naïve, non-adaptive and adaptive groups of mice. C. Graph showing the total (left), mature and stubby (right) dendritic spine density in the hippocampus of naïve, non-adaptive and adaptive groups of mice. D. Representative images of Golgi-Cox stained dendritic segments from the PFC E. Representative images of Golgi-Cox stained dendritic segments from the hippocampus

Following the AST, the brains were removed and Golgi-Cox staining was performed in order to visualize dendritic spines. In the naïve mice (7-8 years old), mature and total dendritic spine density was significantly reduced compared to 3-4 month old adult mice (t-test, p=0.03 for mature and total dendritic spine density) (Figure 5B1). These results further reinforce our results showing the LTP in the PFC was reduced in mice 7-8 months old compared to younger adults (Figure 4A). WM training had a significant effect in the mature (F_(2,12)_=7.1, p=0.01) and total dendritic spine density (F_(2,12)_=8.5, p=0.005), using one-way ANOVA analyses. There was no significant difference among the three groups in the stubby spine density (F_(2,12)_=0.3, p=0.7) (Figure 5B2, B3). In particular, the adaptive groups exhibited increased dendritic spine density to both the non-adaptive and the naïve groups (Tukey’s test, p=0.02 for the naïve vs adaptive and non-adaptive vs adaptive comparisons in the mature and total dendritic spine density). In the hippocampus, WM training had a trend towards a significant effect in mature (F_(2,14)_=4, p=0.03) and no significant effect in the total (one-way ANOVA, F_(2,14)_=2.2, p=0.03) and stubby spine density (one-way ANOVA, F_(2,14)_=0.3, p=0.7). In particular, the adaptive group had significantly increased mature spine density, compared to the naïve and non-adaptive groups (Tukey test, p=0.03) (Figure 5C).

## 4 Discussion

In this study, we find that training mice in the delayed alternation task significantly improves their performance in reversal learning and AST tasks, enhances the LTP in the PFC, increases the fEPSP response in the hippocampus and the dendritic spine density in the PFC. Therefore, WMT enhances cognitive flexibility function as well as functional and structural plasticity, primarily in the PFC.

### 4.1 The role of long-term plasticity and dendritic spines on WM

Because of the short-term nature of WM, studies have focused on short-term physiological and cellular mechanisms that could underlie working memory have been extensively studied in the PFC. Persistent neuronal activity during the delay-period, short-term synaptic plasticity mechanisms and/or neuronal synchronization have been implicated in mediating the ‘on-line’ maintenance and manipulation of information (Mongillo et al., 2008; Riley and Constantinidis, 2015; Miller et al., 2018; Masse et al., 2020). The role of LTP, the cellular correlate of long-term memory, in working memory has not been investigated. However, studies have suggested that even these long-term plasticity mechanisms could be involved in WM tasks, and in particular in learning WM tasks. First, protein synthesis is required for proper performance in working memory tasks (Touzani et al., 2007), which is also required for LTP. While persistent activity is recorded in monkeys before training in working memory tasks (Meyer et al., 2007), the properties of persistent activity change after training (Qi and Constantinidis, 2013; Li et al., 2020). Furthermore, deficits in long-term potentiation correlate with impairments in learning WM tasks (Goto et al., 2009; Brennaman et al., 2011; Konstantoudaki et al., 2017; Chalkiadaki et al., 2019). Therefore, it seems that longer-term plasticity mechanisms are likely to be involved in shaping and/or modulating short-term memory processes, such as working memory.

Dendritic spines are microscopic protrusions from the dendritic tree and represent sites of synaptic contacts of more than 90% of excitatory synapses (Spruston, 2008). Dendritic spine density has been correlated with and predicts WM performance (Hains et al., 2009). In addition, studies have shown that dendritic spine density in the PFC correlates with performance in the AST task (Marquis et al., 2008; Glasper et al., 2015; Stamatakis et al., 2016). Our results further support a significant role for PFC dendritic spine density in learning WM tasks and enhancing cognitive flexibility.

### 4.2 Effect of WM training on HPC function

In our study, we find that WM training, both in the non-adaptive and the adaptive groups, increases the synaptic response to current stimulation and the mature spine density in the hippocampus. These results could also stem from the fact that the alternation procedure is a spatial task, therefore, it engages the hippocampus as well, as evidence by neuronal activity activation during these tasks (Otto and Eichenbaum, 1992). Training in spatial or cognitive control tasks have been shown previously to increase neuronal excitability in the hippocampus (Oh et al., 2009; McKay et al., 2013; Chung et al., 2019). The PFC is interconnected with the hippocampus via different thalamic nuclei (Preston and Eichenbaum, 2013; Varela et al., 2013; Weel et al., 2019), making the possibility for transfer effects from the one brain region to the other plausible.

### 4.3 Effects of WM training on PFC function and cognitive flexibility

Our data strongly indicate that WM training has a beneficial effect on cognitive flexibility as indicated by its effects on reversal learning and AST task. A previous study has shown that WMT (using the eight-arm maze) slightly improves several hippocampal-dependent functions, such as fear conditioning and learning the water-maze task, however, it does significantly improve performance in the mouse-adjusted Stroop task (Light et al., 2010), which also examines cognitive flexibility.

The delayed alternation task is a WM task that depends on PFC function (Sakurai and Sugimoto, 1985; Rossi et al., 2012). Cognitive flexibility depends on PFC sub-regions, depending on the specific task used. The medial PFC is a brain area that underlies WM and participates in changes between attentional sets, while orbitofrontal cortex is necessary for reversal learning (Dalton et al., 2016; Izquierdo et al., 2016). Neuronal firing in medial PFC code for the rule of the task, while neuronal firing in orbitofrontal cortex codes for the response outcome (Simon et al., 2015). The PFC is interconnected directly and indirectly, through the mediodorsal nucleus of the thalamus, with the orbitofrontal cortex (Carmichael and Price, 1996; Alcaraz et al., 2016). Therefore, it is likely that enhanced synaptic plasticity in the medial PFC allow for more accurate neuronal firing in the orbitofrontal cortex and better representations of response outcome, therefore, enhanced facilitation of the formation of new rule response outcome. Our study shows that learning WM tasks increases LTP in the PFC. This enhancement in synaptic plasticity could facilitate information transfer from the medial PFC to other interconnected areas, such as the HPC and the orbitofrontal cortex, allowing the enhancement of cognitive flexiblity. The specific cellular and network events that mediate this transfer are still not understood. However, the results of this study highlight the importance of study the effects of learning and/or cognitive training on the underlying brain areas.

## Acknowledgement

This work was supported by the Dept of Biology, University of Crete and the Special Accounts for Research of the University of Crete (KS) and a Ph.D. scholarship by the Hellenic Foundation for Research and Innovation (HFRI) and the General Secretariat for Research and Technology (GSRT), under the HFRI Phd Fellowship grant (GA. No.4780) (VS).

## References

Alcaraz F, Marchand AR, Courtand G, Coutureau E, Wolff M (2016) Parallel inputs from the mediodorsal thalamus to the prefrontal cortex in the rat. Acsády L, ed. European Journal of Neuroscience 44:1972–1986.

Baeg EH, Kim YB, Huh K, Mook-Jung I, Kim HT, Jung MW (2003) Dynamics of Population Code for Working Memory in the Prefrontal Cortex. Neuron 40:177–188.

Bissonette GB, Powell EM (2012) Reversal learning and attentional set-shifting in mice. Neuropharmacology 62:1168–1174.

Boku S, Izumi T, Abe S, Takahashi T, Nishi A, Nomaru H, Naka Y, Kang G, Nagashima M, Hishimoto A, Enomoto S, Duran-Torres G, Tanigaki K, Zhang J, Ye K, Kato S, Männistö PT, Kobayashi K, Hiroi N (2017) Copy number elevation of 22q11.2 genes arrests the developmental maturation of working memory capacity and adult hippocampal neurogenesis. Mol Psychiatr 23:985–992.

Brennaman LH, Kochlamazashvili G, Stoenica L, Nonneman RJ, Moy SS, Schachner M, Dityatev A, Maness PF (2011) Transgenic mice overexpressing the extracellular domain of NCAM are impaired in working memory and cortical plasticity. Neurobiology of Disease 43:372–378.

Brown VJ, Tait DS (2016) Attentional Set-Shifting Across Species. Current topics in behavioral neurosciences 28:363–395.

Bucur B, Madden DJ (2010) Effects of adult age and blood pressure on executive function and speed of processing. Exp Aging Res 36:153–168.

Carmichael ST, Price JL (1996) Connectional networks within the orbital and medial prefrontal cortex of macaque monkeys. The Journal of Comparative Neurology 371:179–207.

Chalkiadaki K, Velli A, Kyriazidis E, Stavroulaki V, Vouvoutsis V, Chatzaki E, Aivaliotis M, Sidiropoulou K (2019) Development of the MAM model of schizophrenia in mice_ Sex similarities and differences of hippocampal and prefrontal cortical function. Neuropharmacology 144:193–207.

Chung A, Jou CG, Dvorak D, Hussain N, Fenton AA (2019) Learning to learn persistently modifies a neocortical-hippocampal excitatory-inhibitory subcircuit. Biorxiv:817627.

Constantinidis C, Procyk E (2004) The primate working memory networks. Cognitive Affect Behav Neurosci 4:444–465.

Dalton GL, Wang NY, Phillips AG, Floresco SB (2016) Multifaceted Contributions by Different Regions of the Orbitofrontal and Medial Prefrontal Cortex to Probabilistic Reversal Learning. Journal of Neuroscience 36:1996–2006.

Glasper ER, LaMarca EA, Bocarsly ME, Fasolino M, Opendak M, Gould E (2015) Sexual experience enhances cognitive flexibility and dendritic spine density in the medial prefrontal cortex. Neurobiol Learn Mem 125:73–79.

Goldman-Rakic PS (1995) Cellular Basis of Working Memory. Neuron 14:477–485.

Goto Y, Yang CR, Otani S (2009) Functional and Dysfunctional Synaptic Plasticity in Prefrontal Cortex: Roles in Psychiatric Disorders. Biological Psychiatry 67:199–207.

Hains AB, Vu MAT, Maciejewski PK, Dyck CH van, Gottron M, Arnsten AFT (2009) Inhibition of protein kinase C signaling protects prefrontal cortex dendritic spines and cognition from the effects of chronic stress. P Natl Acad Sci Usa 106:17957–17962.

Izquierdo A, Brigman JL, Radke AK, Rudebeck PH, Holmes A (2016) The neural basis of reversal learning: An updated perspective. Neuroscience 345:1–15.

Kamigaki T, Dan Y (2017) Delay activity of specific prefrontal interneuron subtypes modulates memory-guided behavior. Nature Neuroscience 20:854–863.

Konstantoudaki X, Chalkiadaki K, Vasileiou E, Kalemaki K, Karagogeos D, Sidiropoulou K (2017) Prefrontal cortical specific differences in behavior and synaptic plasticity between adolescent and adult mice. Journal of Neurophysiology:jn.00189.2017-39 Available at: http://sci-hub.bz/10.1152/jn.00189.2017.

Li S, Zhou X, Constantinidis C, Qi X-L (2020) Plasticity of Persistent Activity and Its Constraints. Front Neural Circuit 14:15.

Light KR, Kolata S, Wass C, Denman-Brice A, Zagalsky R, Matzel LD (2010) Working memory training promotes general cognitive abilities in genetically heterogeneous mice. Current biology : CB 20:777–782.

Lisman J (2017) Glutamatergic synapses are structurally and biochemically complex because of multiple plasticity processes: long-term potentiation, long-term depression, short-term potentiation and scaling. Philosophical Transactions of the Royal Society B: Biological Sciences 372:20160260.

Liu D, Gu X, Zhu J, Zhang X, Han Z, Yan W, Cheng Q, Hao J, Fan H, Hou R, Chen Z, Chen Y, Li CT (2014) Medial prefrontal activity during delay period contributes to learning of a working memory task. Science 346:458–463.

Lundqvist M, Herman P, Warden MR, Brincat SL, Miller EK (2018) Gamma and beta bursts during working memory readout suggest roles in its volitional control. Nature Communications 9:394.

Lundqvist M, Rose J, Herman P, Brincat SL, Buschman TJ, Miller EK (2016) Gamma and Beta Bursts Underlie Working Memory. Neuron 90:1–14.

Mahmmoud RR, Sase S, Aher YD, Sase A, Gröger M, Mokhtar M, Höger H, Lubec G (2015) Spatial and Working Memory Is Linked to Spine Density and Mushroom Spines. Chapouthier G, ed. PLoS ONE 10:e0139739.

Marquis J-P, Goulet S, Doré FY (2008) Neonatal ventral hippocampus lesions disrupt extra-dimensional shift and alter dendritic spine density in the medial prefrontal cortex of juvenile rats. Neurobiology of Learning and Memory 90:339–346.

Masse NY, Rosen MC, Freedman DJ (2020) Reevaluating the Role of Persistent Neural Activity in Short-Term Memory. Trends Cogn Sci 24:242–258.

McKay BM, Oh MM, Disterhoft JF (2013) Learning increases intrinsic excitability of hippocampal interneurons. The Journal of neuroscience : the official journal of the Society for Neuroscience 33:5499–5506.

Meyer T, Qi XL, Constantinidis C (2007) Persistent Discharges in the Prefrontal Cortex of Monkeys Naive to Working Memory Tasks. Cerebral Cortex 17:i70–i76.

Miller EK, Lundqvist M, Bastos AM (2018) Working Memory 2.0. Neuron 100:463–475.

Mongillo G, Barak O, Tsodyks M (2008) Synaptic Theory of Working Memory. Science 319:1543–1546.

Narayanan NS, Laubach M (2009) Delay activity in rodent frontal cortex during a simple reaction time task. Journal of Neurophysiology 101:2859–2871.

Oh MM, McKay BM, Power JM, Disterhoft JF (2009) Learning-related postburst afterhyperpolarization reduction in CA1 pyramidal neurons is mediated by protein kinase A. Proceedings of the National Academy of Sciences of the United States of America 106:1620–1625.

Otto T, Eichenbaum H (1992) Neuronal Activity in the Hippocampus During Delayed Non-Match to Sample Performance in Rats: Evidence for Hippocampal Processing. Hippocampus 2:323–334.

Preston AR, Eichenbaum H (2013) Interplay of Hippocampus and Prefrontal Cortex in Memory. Current Biology 23:R764–R773.

Qi X-L, Constantinidis C (2013) Neural changes after training to perform cognitive tasks. Behavioural Brain Research 241:235–243.

Riley MR, Constantinidis C (2015) Role of Prefrontal Persistent Activity in Working Memory. Frontiers in Systems Neuroscience 9:181.

Rissman J, Gazzaley A, D’Esposito M (2007) Dynamic Adjustments in Prefrontal, Hippocampal, and Inferior Temporal Interactions with Increasing Visual Working Memory Load. Cereb Cortex 18:1618–1629.

Rossi MA, Hayrapetyan VY, Maimon B, Mak K, Je HS, Yin HH (2012) Prefrontal cortical mechanisms underlying delayed alternation in mice. Journal of Neurophysiology 108:1211–1222.

Sakurai Y, Sugimoto S (1985) Effects of lesions of prefrontal cortex and dorsomedial thalamus on delayed go/no-go alternation in rats. Behavioural Brain Research 17:213–219.

Sala C, Segal M (2014) Dendritic spines: the locus of structural and functional plasticity. Physiological reviews 94:141–188.

Samson RD, Barnes CA (2013) Impact of aging brain circuits on cognition. European J Neurosci 37:1903–1915.

Shoji H, Hagihara H, Takao K, Hattori S, Miyakawa T (2012) T-maze Forced Alternation and Left-right Discrimination Tasks for Assessing Working and Reference Memory in Mice. Journal of Visualized Experiments.

Simon NW, Wood J, Moghaddam B (2015) Action-outcome relationships are represented differently by medial prefrontal and orbitofrontal cortex neurons during action execution. Journal of Neurophysiology 114:3374–3385.

Spellman T, Rigotti M, Ahmari SE, Fusi S, Gogos JA, Gordon JA (2015) Hippocampal– prefrontal input supports spatial encoding in working memory. Nature 522:309–314 Available at: http://sci-hub.tw/10.1038/nature14445.

Spruston N (2008) Pyramidal neurons: dendritic structure and synaptic integration. Nature Reviews Neuroscience 9:206–221.

Stamatakis A, Manatos V, Kalpachidou T, Stylianopoulou F (2016) Exposure to a mildly aversive early life experience leads to prefrontal cortex deficits in the rat. Brain Structure and Function 221:4141–4157.

Tamura M, Spellman TJ, Rosen AM, Gogos JA, Gordon JA (2017) Hippocampal-prefrontal theta-gamma coupling during performance of a spatial working memory task. Nature Communications 8:2182.

Touzani K, Puthanveettil SV, Kandel ER (2007) Consolidation of learning strategies during spatial working memory task requires protein synthesis in the prefrontal cortex. Proceedings of the National Academy of Sciences 104:5632–5637.

Varela C, Kumar S, Yang JY, Wilson MA (2013) Anatomical substrates for direct interactions between hippocampus, medial prefrontal cortex, and the thalamic nucleus reuniens. Brain Structure and Function.

Velázquez-Zamora DA, Garcia-Segura LM, González-Burgos I (2012) Effects of selective estrogen receptor modulators on allocentric working memory performance and on dendritic spines in medial prefrontal cortex pyramidal neurons of ovariectomized rats. Hormones and Behavior 61:512–517.

Velázquez-Zamora DA, González-Ramírez MM, Beas-Zárate C, González-Burgos I (2011) Egocentric working memory impairment and dendritic spine plastic changes in prefrontal neurons after NMDA receptor blockade in rats. Brain Research 1402:101–108.

Wang M, Gamo NJ, Yang Y, Jin LE, Wang X-J, Laubach M, Mazer JA, Lee D, Arnsten AFT (2011) Neuronal basis of age-related working memory decline. Nature:1–5.

Weel MJD der, Griffin AL, Ito HT, Shapiro ML, Witter MP, Vertes RP, Allen TA (2019) The nucleus reuniens of the thalamus sits at the nexus of a hippocampus and medial prefrontal cortex circuit enabling memory and behavior. Learn Mem Cold Spring Harb N Y 26:191–205.

Zaqout S, Kaindl AM (2016) Golgi-Cox Staining Step by Step. Frontiers in Neuroanatomy 10:55–57 Available at: https://www.ncbi.nlm.nih.gov/pmc/articles/PMC4814522/.

